# Molecular identification of whole squids and calamari at fairs and markets in regions of Latin America

**DOI:** 10.1101/2024.05.03.590921

**Authors:** Bianca Lima Paiva, Alan Erik Souza Rodrigues, Igor Oliveira de Freitas Almeida, Manuel Haimovici, Unai Markaida, Patricia Charvet, Vicente Vieira Faria, Bruno B. Batista, Acácio Ribeiro Gomes Tomás, Luis Fernando da Silva Rodrigues-Filho, Jonathan Stuart Ready, João Bráullio de Luna Sales

## Abstract

In recent decades, the commercial importance of cephalopods has increased considerably, being an important fishing resource around the world. However, during the preparation for commercialization of those species, especially those sold in markets, they suffer the process known as *“*calamari*”* which includes removing and separating the head, arm, skin or even having the body structure cut into rings, which ends up making it difficult or often prevents the identification of the species, which can lead to replacements. In this sense, the present study aimed to use the large ribosomal region, rrnL (also known as 16S rDNA) to genetically identify cephalopod species sold in markets and fairs in Latin America. Whole and processed samples were collected from supermarkets and directly from local fishers the approximate collection location. Each generated sequence was submitted to the website *Genbank* for molecular comparison and included in the database for subsequent genetic identification. Comparison of sequences using the *Blastn* revealed the presence of eight species that are widely traded in the Latin American region. Our results indicate labeling errors in samples from the State of Pará that contained the species *Dosidicus gigas* (d’ Orbigny, 1823) found only in the Pacific Ocean and were generically labeled as “National Lula”. No type of substitution was found among the samples that were being sold at fairs and markets, only labeling errors. Thus, our results demonstrate the effectiveness of the rrnL for identifying species and evaluating labeling errors.

## Introduction

In recent decades, the commercial importance of cephalopods has grown notably in volume of trade (Caddy and Rodhouse 1998; Watson and Pauly 2001; Jereb and Roper, 2010). The number of cephalopod species that are targets of commercial fisheries has grown in recent times (FAO 2022), and given this, they represent the third largest marine group consumed by the human population (the largest two being finfish and crustaceans) (Gleadall et al 2024). And when it comes to commercial value, members of the family Ommastrephidae represent the largest volume of commercialization and global importance (FAO 2016). Loliginids represent the second family of interest for global captures due to their distribution limited to the continental shelf, in tropical and temperate regions around the world, as well as the high quality of the species’ meat (Jereb and Roper 2010; Sales et al. 2011), having a high protein value, a wide variety of essential amino acids and extremely low fat levels, making regular consumption of the species part of healthy and balanced diets (Anfaco-Cecopesca 2018; USDA 2018).

Currently, cephalopods represent about 7% of seafood production, their landings have increased since 1961, reaching a record of captured and commercialized tons in 2020 (FAO, 2022). In terms of cephalopods catches and consumption, Asian, European, African and North American countries have the ten largest fishing fleets in the world (FAO 2020), with Spain, Italy and Japan being the largest consumers (FAO 2016; 2018).

In South America, *Dosidicus gigas* D’orbigny 1835 has been widely explored, being exported by Peru to more than 50 countries, where several attempts are being made to diversify new products made with the species among them, canned, pre-cooked, roasted and others (Vieites et al. 2019). This high demand for export means that the commercialization of entire species is greatly reduced or made difficult, with the need for processing often resulting in a longer shelf life of derived products (Anfaco-Copesca 2018; USDA 2018).

However, processing most often removes or damages diagnostic features that are important for correct species identification through traditional taxonomic characteristics (Di Pinto et al. 2013). In the case of cephalopods, there is the removal of the intestines, skin, arms, tentacles, fins and head, where the species are often cut into rings or tubes (Chapela et al 2003; Johnson 2007), thus removing all morphological characteristics which are used to identify species as sex, or sexual maturity, a process that make all species be commercialized as calamari (Roper and Mangold 1998; Chapela et al. 2003).

These modifications make taxonomic identification difficult or even impossible, making it more prone to economic fraud where highly valued species, such as those from the family Loliginidae, are replaced by species of lower commercial value, such as those from the Ommastrephidae, which becomes a concern for the international commercial market (Johnson 2007; Chapela et al. 2003), or often, unintentional substitution, where due to lack of systematic knowledge, suppliers themselves are unable to correctly identify the species, assigning them umbrella terms that shelter many different species (Cawthorn et al. 2013; Kroetz et al. 2020; Sharrad et al. 2023).

Molecular methods have been widely used, firstly, to identify new species or populations of different species, thus providing a complement to pure systematic studies, with their applicability in the identification of species already consolidated in several different taxonomic groups (Virgilio et al. 2010; Hollingsworth et al. 2011, Gales et al. 2022; Rodrigues-Filho et al. 2023), especially DNA barcode methods (Hebert et al. 2003). This technique is based on analyzing the variability of a short nucleotide sequence to assess differences between species (Hebert et al. 2003). The cytochrome oxidase subunit I (*coxI*) gene was initially proposed by Hebert et al. (2003), this region of the mitogenome includes both primary and conserved sites, as well as a suitable sequence of variation that allows differentiation between species (Hellberg et al. 2011).

There have been many criticisms regarding the choice of the *cox1* gene as a barcode region, such as the presence of pseudogenes (Rubinoff 2006; Rubinoff et al. 2006). For mollusk species this is a major problem since several studies have already demonstrated the presence of alterations in the mitochondrial genome, including reorganizations, gene duplications and deletions (Boore and Brown 1995; Boore 1999), in addition to bi-parental inheritance in bivalve species (Hoeh et al. 1996). For cephalopod species, the main limitations of the application of the *cox1* gene are found in the Ommastrephidae family due to several species having two copies of this gene (Yokobori et al. 2004), making its application as a DNA barcoding tool inappropriate for this group. The large ribosomal region rrnL (also known as 16S rDNA) is a suitable alternative to be used as a molecular region for species identification, due to its conserved nature and large number of copies in mitochondria, having already been widely used in resolving phylogenetic analysis in cephalopod species (Bonnaud et al. 1994; Warnke et al. 2004; Sales et al. 2013; Sales et al. 2019; Costa et al. 2021).

Among the main advantages of using molecular methods in the identification of fishery products is the fact that it is easy to amplify target regions, as well as the fact that any portion of the body of a specimen or individuals from preparation processes such as cooking, frying, canning can be used (Teletchea et al. 2005; Villanueva-Zaya et al. 2021; Mottola et al. 2022). In this way, the applicability of molecular methods can help in the correct identification of the species being commercialized, which can reveal not only economic losses due to substitutions, as well as identify commercialization’s of species that may pose health risks (Von der Hayden et al. 2010) or the presence of endangered species with prohibited capture, fishing and landing status, making it necessary to use appropriate tools to confirm product labeling and avoid commercial fraud (Barbuto et al. 2010). Reliable molecular tools for barcoding DNA analysis have been developed to protect consumers from food frauds and health hazards and to improve the monitoring of endangered species due to overfishing and illegal commercial activities (Filonzi et al. 2023).

When complemented by molecular methods, market research can provide accurate species identification, even for products that have been through the finning and processing before sale (Palumbi 2007). DNA methods have been widely used for forensic analysis of food products because they provide an efficient, informative, sensitive and specific identification stimulus and because they can be applied to highly processed food products (Hebert et al. 2003; Rasmussen and Morrissey 2008).

In Brazil, legislation on the cephalopod trade does not require all products to bear the common name and scientific name of the species marketed on their labels, which can lead to the intentional substitution of fish-based products, which can cause economic loss to consumers. The European Union, in an attempt to seek solutions to reduce commercial fraud, has instituted regulations (EC 104/200; EC 2065/2001) that require member countries to make official lists available with scientific, common and commercial names of fish species that are being marketed, as well as requiring that product labels also be clearly informed (Brito et al. 2015).

Taking into account all the problems exposed regarding the commercialization of cephalopods, as well as the absence of official legislation labeling these products that are sold in Brazil. The present study aimed to use the fragment of the rrnL to identify which species of squid are being sold, both in the form of trays containing processed animals, and specimens sold whole in markets and fish fairs in some Latin American countries investigating whether there is evidence of intentional substitutions in the trade of cephalopod species.

## Material & Methods

### Sampling

For the present study, both whole and processed squid were collected directly from fishers or from supermarket chains in different regions of Brazil, Suriname, Mexico and French Guiana. For some samples, the place of origin (fishing) and purchase was different, with all information relating to these cases also duly recorded (samples purchased in Fortaleza-Ceara state, Brazil, had identification of origin, coming from Argentina and another package purchased in Belém-Para State, Brazil, also identified the origin of the content as Chile, Figure 1). Subsequently, a small piece of muscle tissue was removed from each sample stored in 70% ethanol and subsequently stored at -4°C until DNA extraction. About samples purchased in a processed form, each ring contained in the packaging was considered as a different sample. Each product tag with the assigned generic identification name, as well as the name of the distributor and fishing company were registered.

**Fig. 1:**
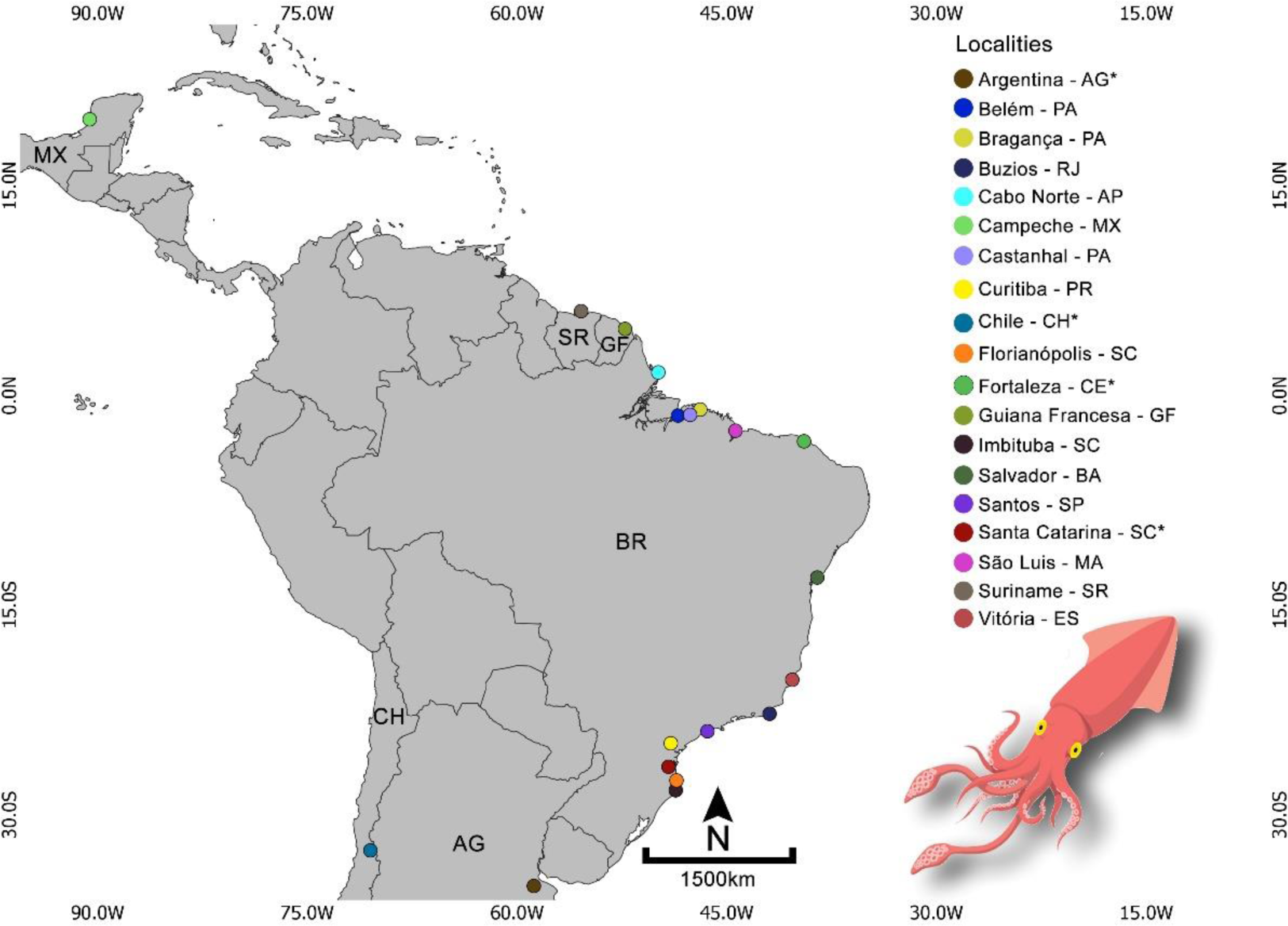
Map showing all squid sampling places utilized in the present study. Colored circles correspond to each localities. See Table 1 for further definitions for localities names marked with *. Abbreviations represent the capitals name of each countries where: AG=Argentina; BR=Brazil; CH=Chile; GF=Guiana Francesa; MX=Mexico; SR=Suriname. Map created with Ogis. Shapefile with projection WGS84. All Editions were made on Photoshop Cc2023. Source: Shapefile IBGE.

### DNA Extraction, PCR and Sequencing

Total DNA was extracted from muscle tissue using the CTAB DNA extraction protocol (Doyle and Doyle 1987). The rrnL fragment was isolated and amplified by polymerase chain reaction (PCR) using the primers: L1987 (5’-GCCTCGCCTGTTTACCAAAAAC-3’) and H2609 (5’-CGGTCTGAACTCAGATCACGT-3’) (Palumbi et al. 1991). The reactions were carried out containing a final volume of 15 µl, with 2.4 µl of dNTP (1.25mM), 1.5 µl of buffer solution (10X), 0.8 µl of MgCl_2_ (50 mM), 0.6 µl of each primer, 0.12 µl of Taq DNA polymerase (5 U/ul), 1 µl of genomic DNA (200 ng/ul), and ultrapure water to complete the final reaction volume.

Reactions were performed according to Sales et al. (2013), whose protocol consists of: initial denaturation at 94°C for 2 min, followed by 30 cycles of denaturation at 94°C for 30 s, hybridization at 51°C for 1 min and extension at 72°C for 2 min, with the final extension step of 72°C in a period of 7 min. The quality and size of the PCR product were checked using 2% agarose gel electrophoresis. For sequencing of the fragments obtained, the PCRs were previously purified using 65% isopraponol and 70% ethanol, and the sequencing reactions were carried out with reagents from the BigDye Terminator V3.1 Cycle Kit and then sequenced on the SeqStudio Genetic Analyzer sequencer (Applied Biosystems).

### Data analysis

The generated DNA sequences were aligned with the CRUSTAL W automatic alignment tool (Thompson et al. 1997) implemented in the Bioedit v7.0.9.1 program (Hall 1999). Subsequently, the sequences underwent visual inspection for possible automatic alignment errors, mainly in the hyper variable region of the rrnL region.

Two validation methods for molecular identification of the samples were used in the present study. At first, the *Blastn* tool, available on the GenBank portal (https://www.ncbi.nlm.nih.gov/genbank/) was used following a genetic similarity criterion of 98% for the taxonomic signature specific to each sequence. Based on the result of *Blastn*, sequences available from NCBI were aggregated for subsequent phylogenetic inferences with the following species: *Doryteuthis pleii* Blainville, 1823; *Doryteuthis pealeii* Lesueur, 1821; *Doryteuthis sanpaulensis* Brakoniecki, 1984; *Lolliguncula brevis* Blainville, 1823; *Illex argentinus* Castellanos, 1960; *Dosidicus gigas* d’ Orbigny, 1835; *Nototodarus sloanii* Gray, 1849; *Uroteuthis duvaucelii* d’ Orbiny, 1835. The number of sequences generated in this study with the access code, as well as the sequences downloaded from GenBank are available in supplementary material 1. Additionally, for species delimitation as a second identification method, the TIM+F+G4 evolutionary model was estimated by ModelFinder software (Kalyaanamoorthy et al. 2017), as the best model to our dataset. Subsequently, a Maximum Likelihood phylogenetic tree was estimated with IQ-tree v2.2.0 program package (Minh et al. 2020) using Ultrafast Bootstrap (Hoang et al. 2018) based on 1000 pseudoreplicates. The software Figtree v1.4.4 (Rambaut 2010) was used to edit the phylogenetic tree. To visualize the data, Barplot and PieDonut were generated in Software R (R Core Team 2022), using the “ggplot2” and “webr” packages, respectively.

## Results

### Molecular identification

A total of 552bp of the rrnL mitochondrial gene fragment were generated from 181 squid samples. Comparison of sequences using the *Blastn* revealed the presence of eight species that were widely traded throughout the Latin American region, namely: *D. pleii, D. pealeii, D. sanpaulensis, L. brevis, I. argentinus, D. gigas, N. sloanii, N. gouldi and U. duvalcelii* (Table 1).

**Table 1.** Squid species identified in the present study based on rrnL fragment (16S rDNA). Data about sampling places, label identification, kind of processing, and genetic similarity match is provided. Numbers above some of the sampling places indicate the actual origin place from where the samples came: 1 – Santa Catarina State; 2 – Argentina; 3 – Chile. See Supplementary Material 1 for detailed table showing the *Blastn* results per sample.

All sequences evaluated in the present study, when compared to data from Genbank, obtained an identification at species level with >98% similarity. Of the 181 samples identified, 18 samples showed 100% similarity with *D. pleii* (n=13), *D. pealeii* (n=2) e *L. brevis* (n=3), from the following locations respectively: Bragança, Florianópolis, Búzios, Campeche and Curitiba. The remaining of the identified samples (47) had a minimum similarity value of 99.80% with *D. pleii* (n=9), *L. brevis* (n=28), *D. sanpaulensis* (n=5) e *D. pealeii* (n=5), from the following locations: São Luís, Imbituba, Santos, Cabo Norte (all four from Brazil), and Suriname.

Furthermore, 11 samples showed a minimum similarity value of 99.79% with *I. argentinus* (n=5) *and D*. *pleii* (n=6) coming from Santa Catarina, Argentina and Suriname. 34 samples showed a similarity value of 99. 59% with the species *D. pleii* (n=8) and *D. pealeii* (n=26) from Salvador, Fortaleza, Vitória and Cabo Norte, respectively. Among the samples from French Guiana and Belém, 11 samples showed a minimum similarity value of 99.43% with the following species, *U. duvaucelii* (n=6)*, D. pleii* (n=4) and *D. gigas* (n=1). Finally, 36 samples obtained a similarity value of 99.41% for *D. gigas*, from Chile (n=9) and Castanhal (n=27). Eight identified samples showed a similarity of 99.39% with *D. pleii*, coming from Cabo Norte. Among the samples from Florianopolis, six showed a minimum similarity value of 99.21% with *I. argentinus* while 10 remaining samples from French Guiana showed a value of 98.88% for *N. sloanii.* No kind of species substitution (intentional or not) was found in the present study.

Twenty-nine sequences were added from GenBank, based on the degree of similarity of previously identified sequences, 27 of which are related to species identified with *Blastn* and three sequences of the species *N. gouldi*, resulting in a final alignment of 210 sequences. All sequences previously identified in *Blastn* were recovered in highly supported clades, separating two genetically distinct families, Loliginidae and Ommastrephidae, with five and three species identified, respectively. The results found with the Maximum Likelihood tree were in agreement with the genetic similarity analysis (NCBI-Blast), except in some cases referring to the structures within some species (Fig. 2).

**Fig 2.**
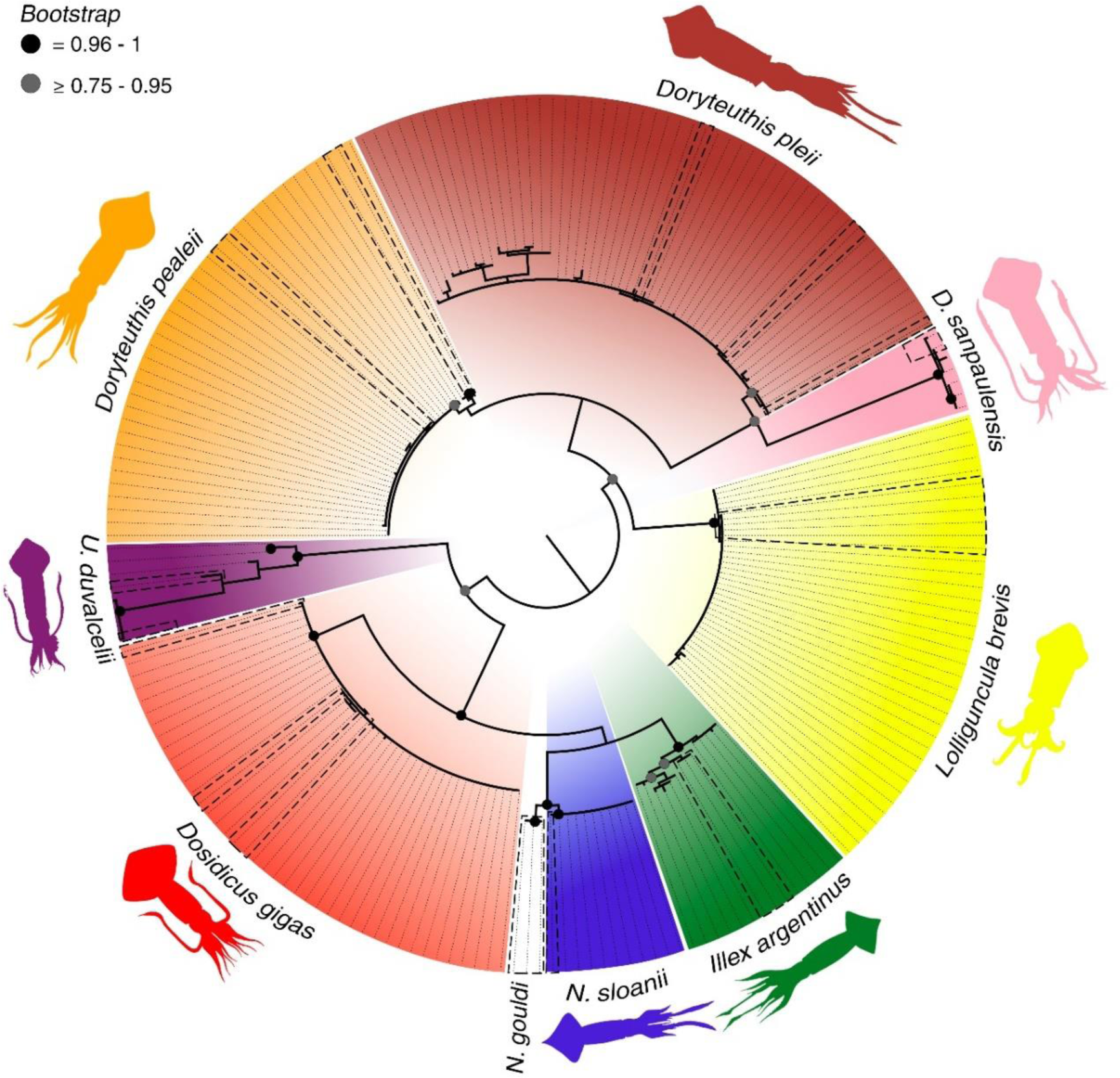
Maximum Likelihood phylogenetic tree of the squid species identified in the present study based on rrnL mitochondrial gene fragment (16S rDNA). Each species is represented by its respective colored outline illustration: *D. pleii* – Brown; *D. pealeii* – Orange; *D. sanpaulensis* – Pink; *L. brevis* – Yellow; *D. gigas* – Red; *U. duvalcelii* – Purple; *N. sloanii* – Blue e *I. argentinus* – Green. Dashed shapes indicate the positioning of DNA sequences obtained from GenBank that were associated to each respective species clade. Image processing was done using the Adobe Photoshop Cc2023 software.

Within the family Loliginidae, three genera were recovered, *Lolliguncula*, *Doryteuthis* and *Uroteuthis*, totalizing 123 individuals identified. The most abundant species was *D. pleii* with 48 individuals forming a branch with 78% bootstrap support associated with different regions (Table 1, Figure 2). *Dorytheuthis pealeii* corresponds to the taxon with the second highest abundance among Loliginidae, totaling 33 identified sequences forming two clades with support values of 96% and 78%, respectively. *Lolliguncula brevis* (n=31) formed a group with the sequences downloaded from Genbank according to the respective taxon with a support value of 97%. *Urotheutis duvaucelii* (n= 6) was recovered with a high support value (99%), however there is evidence of strong distinct grouping structures with sequences from GenBank. *Dorytheuthis sanpaulensis* (n=5) formed a cluster with 100% support value with a structure established between sequences from Santos-SP.

Species from three other genera were also identified, *Dosidicus*, *Illex* and *Nototodarus* (all belonging to the family Ommastrephidae), with a total of 58 individuals. The most abundant species for this family was *D. gigas* with 37 individuals forming a branch with 99% support value. *Illex argentinus* (n=11) formed a clade with 98% support with internal structuring being recovered with 87% support from sequences belonging to samples from Florianopolis In the case of sequences of *N. sloanii* (n=10), 97% support value separated them from *N. gouldi*, also demonstrating a support value of 99% that highlights this difference between the sequences, since the grouping with *N. sloanii* got the value of 97% together with the sequences from Genbank. In the previous analysis, some sequences obtained high genetic similarity with *N. gouldi*, but it was possible to observe the separation with a high bootstrap value among the congeners present in the database.

### Squid Processing level

Most of squid sampled (71%) were being sold in whole form, while the remaining in processed form (29%) (Fig. 3a). The majority of individuals sold comprise Loliginidae, especially individuals that were being sold (whole) in their entirety at fairs. The most common species found was *D. pleii* (Fig. 3b) from the Southwestern Atlantic distributed among the states of Amapá, Pará, Maranhão, Bahia, Rio de Janeiro, Espírito Santo and Santa Catarina, all samples were being sold whole in supermarkets and fairs.

**Fig 3.**
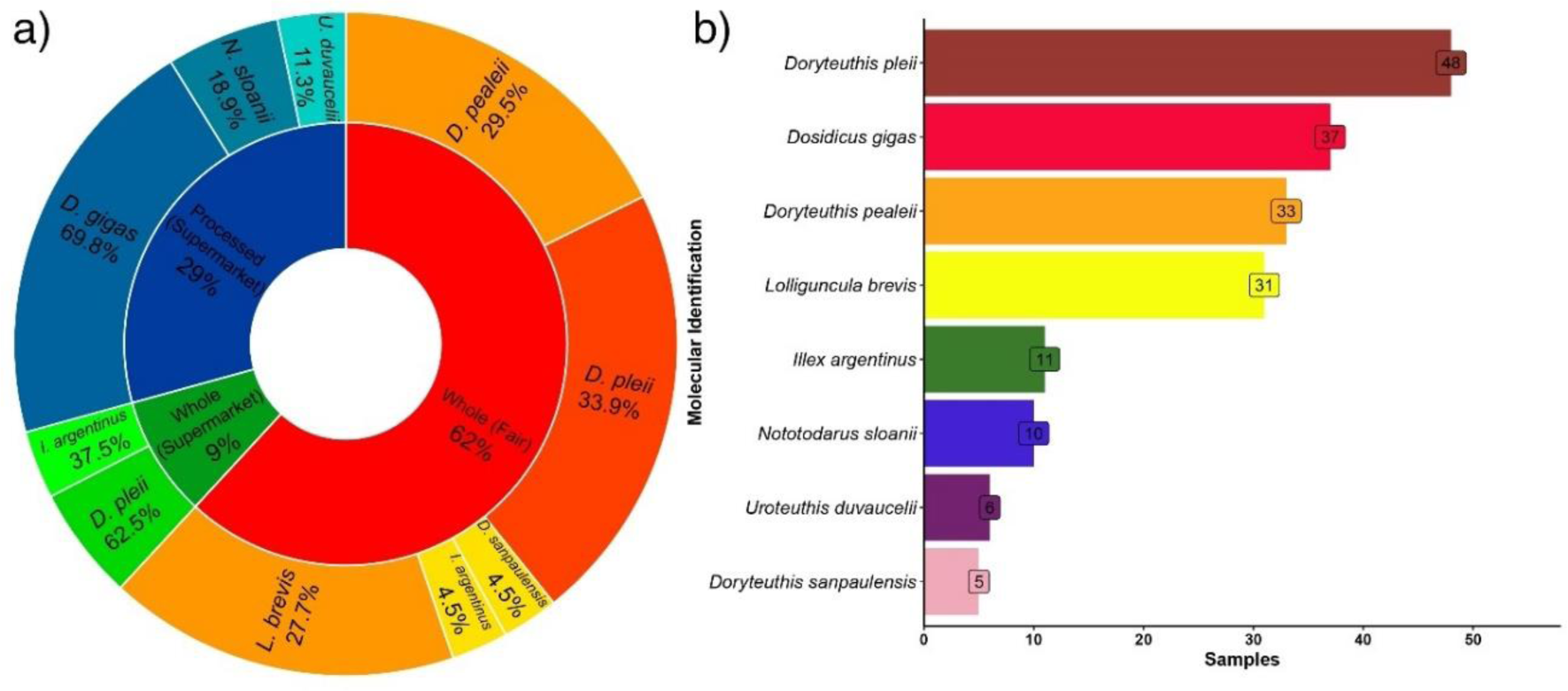
Graphs demonstrating the quantified results of the present study. a) PieDonut indicating the samples identified based on the type of processing and place of commercialization of the squid. b) Barplot summarizing the abundance of commercialized squid species.

The second most frequently found species was *D. gigas* purchased in the State of Pará. In the case of this species, the individuals were being sold in a processed form (cut into rings), containing generic information on the labels as “*Lula Nacional*” (National Squid), “*Lula Brasileira*” (Brazilian Squid), or just “*Lula*” (Squid). The occurrence of labeling errors was observed in 100% of the products, when these samples come from the cities of Belém and Castanhal. Only one sample that was being sold in a supermarket in Belem was being sold without any information on the product label, containing only on the supermarket freezer label such as “Squid in rings”. The third most common species was *D. pealeii* from Cabo Norte (Brazil) and Mexico. Another common species was *L. brevis*, being present in fairs from two locations Cabo Norte and Santos. All samples identified in the present was commercialized as whole being sold in fairs.

Among the less frequently traded species identified in this study were *I. argentinus*, found both in fish markets in the city of Fortaleza but with Argentina as the place of origin on the label, and supermarket chains in Santa Catarina as a whole. The species *N. sloanii* was also identified less frequently, being sold in French Guiana in processed form (salads) in supermarkets, followed by *U. duvaucelii* which was also sold in French Guiana in the same way of *N. sloanii*, followed by *D. sanpaulensis* and for *D. pealeii* from the Equatorial Northwest Atlantic, a genetically distinct lineage of *D. pealeii* from North Atlantic (Sales et al. 2024).

## Discussion

Incorrect labels end up deceiving OU jeopardizing consumers’ conscious choices, posing potential health risks (Tartre 2016), as well as generating financial losses for consumers (von der Heyden et al. 2010). Additionally, from a conservation point of view, they “mask” the commercialization of species with prohibited fishing status (Palmeira et al. 2013; Zeng et al. 2019) or, when commercialized throughout the year, species that are undergoing period of reproductive closure end up being captured, thus causing possible future impacts on their recruitment and population stock (Stawitz et al. 2016).

Processing often removes or damages diagnostic characteristics that are important for the identification of species through conventional taxonomy (Barbuto et al. 2010; Di Pinto et al. 2013), which ends up contributing to the possible replacement of species of greater value commercial for others of lower value. The results generated in the present study indicate that a number of labeling errors occurred in products that had specimens sold in 2015 in a processed form as squid rings sold with the generic name “Lula Nacional” (National Squid), a name that suggests to the consumer that the product was caught in Brazil. All samples derived from squid labeled this way were genetically identified as *D. gigas,* a squid species found exclusively in the Pacific Ocean (Markaida et al. 2005; Jereb and Roper 2010). A single sample that was being sold in processed form at the supermarket in Belém, with identification of “*Lulas em anéis*" (Squid Rings) in June 2022 which was also identified as *D. gigas*, was not in accordance with SDA Ordinance No. 544, of March 14, 2022 (article 16 in paragraph 1), where it is emphasized that “For products of the type *D. gigas,* it is mandatory to include scientific nomenclature, such as the explanatory affix to the product’s sales name”. This is the only species of cephalopods for which there is an ordinance that regulates commercialization in all Brazilian territory.

Nevertheless, the replacement of commercial species cannot always be characterized as intentional commercial fraud. A portion of inappropriate labeling occurs unintentionally, largely due to the morphological characters used to identify species that can be easily mistaken, or simply when using vernacular names that are common to more than one species. To a certain extent, mislabeling may be the result of difficulties in identifying species, which occurs mainly in groups with few informative morphological characters and with a long history of cryptic species, such as in the Loliginidae species (Sales et al. 2013; Cheng et al. 2014; Costa et al. 2021, Sales et al. 2024a) and Ommastrephidae (Fernández-Álvarez et al. 2020). Confusion can also arise due to different species having the same vernacular name, or different common names in different regions for the same species (Brito et al. 2015). However, usually the substitutions of species, which always characterize commercial fraud, aims to obtaining higher profits (Pauly et al. 2005; Worm et al. 2006).

Regarding the species purchased whole from markets in the Equatorial Northwest Atlantic that were identified in the present study, some considerations must be made. The squid fauna proposed for these regions is very similar to that reported for some regions of the North Coast of Brazil (Jereb and Roper, 2010). The presence of *D. pleii*, *L. brevis* and *D. pealeii* in these locations are in accordance with what has been proposed by recent molecular reviews with specimens from the North Coast of Brazil (Sales et al. 2011; 2013; 2014; 2017; 2024a; Costa et al. 2021). Two species of *Doryteuthis* identified in the present study are of high commercial importance in Brazil and in North America (Herke and Foltz 2002; Jereb and Roper 2010; Postuma and Gasalla 2015), and, it is important to clarify that the lineage of *D. pleii* commercialized in Brazil corresponds to the actual species (Sales et al. 2017), while the lineage exploited in North America corresponds to a not yet described species. The same specific delimitation is valid for *L. brevis* (Sales et al. 2014) and, although this is not a commercially important species (Good et al. 2023), on the North coast of Brazil it is often consumed by fishers (Sales et al 2024b). For *D. pealeii*, the pattern is the opposite, with the lineage that occurs in Brazil being a new undescribed species (Sales et al 2024a). According to our results, most of the samples were in agreement with the fauna found on the Brazilian coast (Haimovici et al. 2009), indicating that the species that were being sold at fairs come from artisanal fishing, which contributes to the lack of supervision by the responsible governmental authorities, making it difficult to verify frauds. The lack of evidence of substitution among the products that were used in the present study denotes the difficulty in verifying the occurrence of substitution in samples that are sold in Brazil, due to the lack of information on the label, such as the name of the specific species name being sold for the consumer, thus making it difficult to track the replacement of one species of squid by another of the same genus, or family.

For the species in the present study, in relation to other species sold in markets in Brazil and in Latin American countries utilized here, there was no case of commercial fraud, but rather a labeling error, since the specimens were sold only as “Squid”. The sequences that were identified as *U. duvaucelli* demonstrated evidence of strong different clades with sequences originating from the GenBank. This result is in agreement with other previous studies that already indicated the presence of multiple lineages within *U. duvaucelli*.

*U. duvaucelii*, presents a relevant morphological and genetic polymorphism within a wider geographic area (Sin et al. 2009), and is of commercial importance in the Asian region, being mainly fished by artisanal fleets, but it already constitutes a high volume of exports, specially from Thailand to other countries. This species represents 68% of the cephalopod species exported by India (Jereb and Roper 2010). The clade formed in the Maximum Likelihood (ML) tree reinforces this finding, although, the identification with *Blastn* suggests that all sequences generated in the present study correspond to the taxon in question. Different morphotypes of *U. duvaucelii* are recognized by commercial fishing. One of these are formed by a relatively large and a smaller form from the Gulf of Aden and the Arabian Sea (Nesis 1987), and two other forms are known, one more robust and one more slender from the Eastern Pacific (Okutani 2005). In this sense, the diversity of the group may be underestimated, significantly affecting the management and conservation of the species. The results of molecular reviews carried out in recent years reinforce the presence of different species within *U. duvaucelli* (Sales et al. 2013; Krishnan et al. 2022). One of the bottlenecks from the point of view of fisheries management and resource exploitation arises precisely when a given species targeted for exploitation has a cryptic species, because from the point of view of management and commercial exploitation, it would not be a single species under exploitation, but two or multiple at the same time, which can lead to a reduction in the population stock of these lineages, as the reproductive period and breeding area are often different between them (Avise 1996).

Finally, the sequences of *N. sloanii* were well separated in the phylogenetic tree, although, for some sequences *Blastn* indicated a high similarity with *N. gouldi.* However, in the same topology, *N. sloanii* and *N. gouldi* formed distinct clades with high support value. One possible cause for this is the high level of similarity between species since until recently *N. gouldi* was considered a subspecies of *N. sloanii.* According to the information contained on the packaging label that these samples came from FAO fishing area 81 (Southwest Pacific), which is consistent with the species’ distribution area. It is worth noting that the samples identified as *N. sloanii* came from French Guiana and were being sold processed in supermarkets, containing all the information regarding the labeling described by the European Union in the “EC 104/200” regulation; EC 2065/2001”. Data obtained from FAO indicate that global catches for this species decreased in 2021, reaching a volume of 29,832 tons, data lower than the years 2020 and 2019, which had a volume of 41,928 and 43,795 tons, respectively.

As there is no comprehensive Brazilian legislation for cephalopods that requires the common and scientific name of all species marketed labels, there is also no official list containing the common, commercial, or scientific names of the species sold in the country as well for Latin America Countries.

## Conclusion

Based on the results of the present study, the samples identified as *D. gigas* that were being sold in a processed form (rings) in supermarkets, were considered incorrect labeling. For all other taxa identified in the present study, no occurrence of intentional or unintentional replacement was found. Due to the fact that cephalopod species present high morphological plasticity, the use of molecular tools becomes necessary for the correct identification and labeling of species that are commercialized. With the high levels of incorrect labeling recorded, the present study shows the importance of using molecular methods that help in the identification of squid species that are commercialized. The lack of specific Brazilian legislation on labeling for all Brazilian fauna cephalopod (as such Latin America Countries) products may favor intentional and unintentional substitutions, bringing problems such as economic fraud or species substitutions. We are also relieved that our results continue to debunk the urban myth of calamari substitutions with pig rectum (Grenoble, 2013).

## Acknowledgments

The Instituto Tecnológico Vale-Desenvolvimento Sustentável (ITV-DS) provided laboratory infrastructure to carry out the DNA sequencing and we thank Dr. Silvia Britto Barreto for supporting the laboratory procedures.

## Statements and Declarations

### Ethics approval

The research was developed under the ethical guidelines of Universidade Federal do Pará.

### Consent to participate

Authors declare consent to participate.

### Consent for publication

Authors declare consent for publication if the manuscript is accepted.

### Author Contributions

Conceptualization, J.B.L.S and B.L.P.; Formal analysis, B.L.P., I.O.F.A. and A.E.S.R.; Methodology, M.H., U.M., P.C., V.V.F., B.B.B., A.R.G.T., and L.F.S.R-F.; Writing—review and editing, B.L.P., I.O.F.A., A.E.S.R., M.H., U.M., P.C., V.V.F., B.B.B., A.R.G.T., L.F.S.R-F., J.S.R., and J.B.L.S;

### Data Availability Statement

The sequence data are in submission processes at the GenBank repository;

## Supplementary Material

Supplementary material 1: Metadata of the sequences used in the present study. In bold, they indicate sequences downloaded from NCBI-Genbank.

Supplementary material 2: Maximum Likelihood (ML) phylogenetic tree with support values.

Supplemental material 3: A) R Scrypt built to generate PieDonut. B) Datasheet used to generate the graph.

Supplemental material4: A) R scrypt built to generate the bar chart. B) Datasheet used to generate the Barplot.

