## Supplementary figures and images for "Molecular identification of whole squids and calamari at fairs and markets in regions of Latin America"

### Supplementary file 2

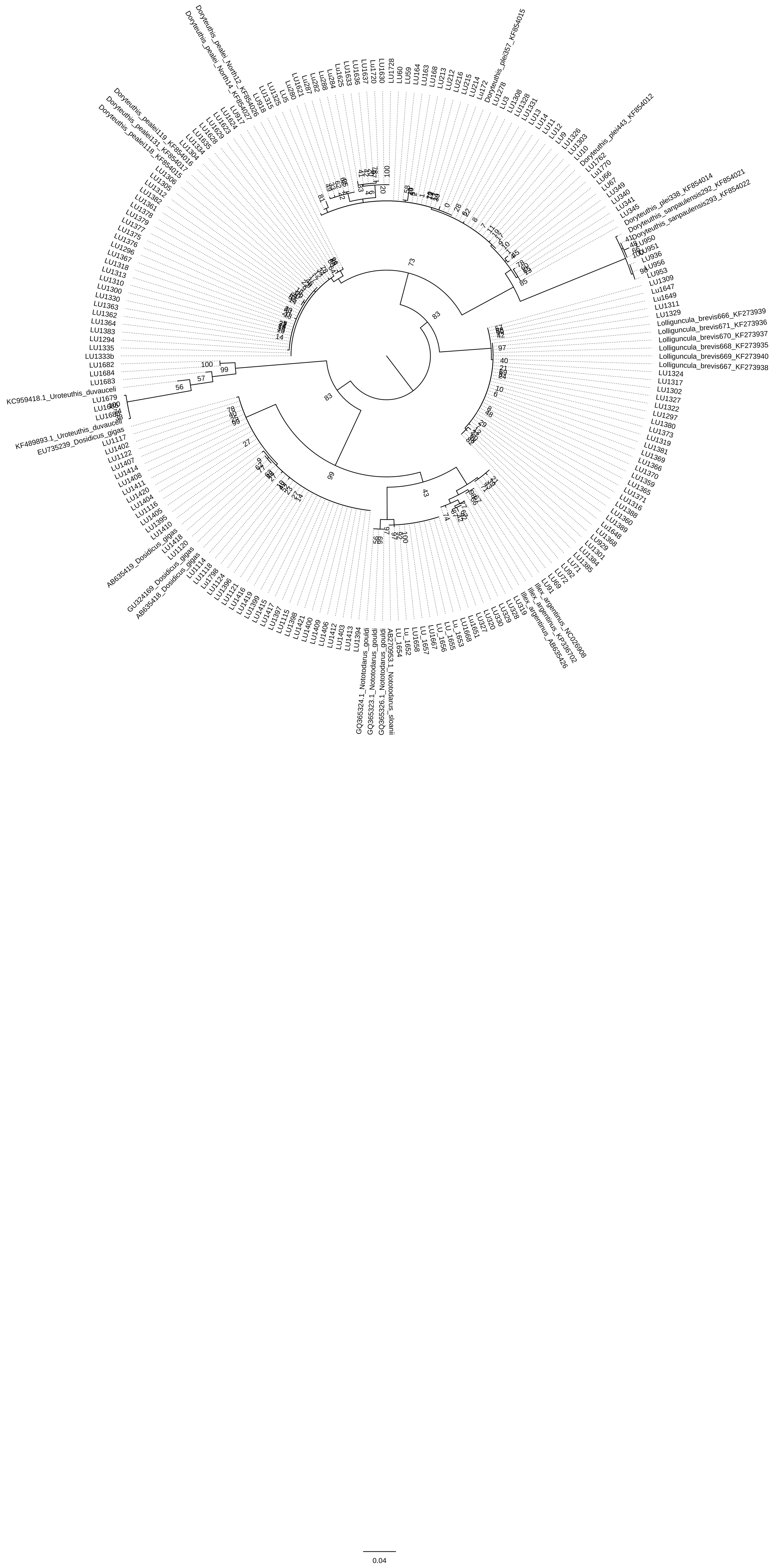
